# Towards a bioelectronic computer: A theoretical study of a multi-layer biomolecular computing system that can process electronic inputs

**DOI:** 10.1101/290775

**Authors:** Katherine E. Dunn, Martin A. Trefzer, Steven Johnson, Andy M. Tyrrell

## Abstract

DNA molecular machines have great potential for use in computing systems. Since Adleman originally introduced the concept of DNA computing through his use of DNA strands to solve a Hamiltonian path problem, a range of DNA-based computing elements have been developed, including logic gates, neural networks, finite state machines (FSMs) and non-deterministic universal Turing machines. It has also been established that DNA molecular machines can be controlled using electrical signals and that the state of DNA nanodevices can be measured using an electrochemical readout. However, to the best of our knowledge there has as yet been no demonstration of a fully integrated biomolecular computing system that has multiple levels of information processing capacity, can accept electronic inputs and is capable of independent operation. In this paper we address the question of how such a system could work. We present simulation results showing that such an integrated hybrid system could convert electrical impulses into biomolecular signals, perform logical operations and take a decision, storing its history. We also illustrate theoretically how such a system could potentially be used to perform a task such as controlling an autonomous robot navigating through a maze.

## INTRODUCTION

Adleman demonstrated in 1994 that a system of DNA molecules could solve the travelling salesman problem^1^ and since then great progress has been made in the field of DNA computation. A wide variety of DNA logic gates have been designed, with operating mechanisms including strand displacement^2,3^ and DNAzyme action^4^. It has been shown that ‘see-saw’ gates^5^ can be used for thresholding and the construction of complex DNA computing systems that are capable of performing arithmetic functions^6^ or demonstrating memory and inference^7^. Other DNA computing paradigms include chemical reaction networks^8^, finite state machines^9,10^ or algorithmic self-assembly of tiles^11^, and it was recently demonstrated that DNA could be used to implement a non-deterministic universal Turing machine^12^. The capabilities of DNA computing machines could be broadened further via their integration with electronic systems, which could underpin new approaches for input, processing and readout^13^.

It has already been demonstrated that DNA molecular machines can be controlled using electrical signals, using electrochemical control of pH and pH-sensitive structures such as i-motifs or triplexes^14,15^ or electronically triggered release of a signalling species from a surface^16^. In an alternative architecture, an enzymatic logic gate under certain conditions produces NADH, which activates an electrochemical process that leads to release of DNA from an alginate matrix^17^. For these studies, electrical triggers were used to supply inputs, but electrochemical effects can also be used for signal readout. For instance, E-DNA sensors consist of DNA nanoswitches functionalized with a redox active moiety that can transport charge to the surface of an electrode when the structure is in one configuration but not when it is in an alternative state^18^. It is also possible to use the electrochemical signal from intercalated electroactive molecules (such as methylene blue) to confirm whether DNA is single-stranded or hydrogen-bonded in a duplex, quadruplex etc^19^.

In most cases, these electrochemical systems have a single layer of information processing capacity. In other words, they simply yield a particular output in the presence of a stimulus. While such devices are extremely useful for biosensing applications, more complex systems are required for computing. In this paper we present a proof-of-concept theoretical study of a new type of biomolecular computer. Our goal was to design and theoretically validate a system with multiple layers of information processing capacity, based primarily on well-characterized DNA machinery, with the inputs being provided electronically. We incorporate a diverse range of DNA elements, from pH-sensitive nanoswitches to catalytic cycles and logic gates. Our simulations focus on the kinetics of the DNA machines, and as each operating cycle of our DNA-based computer takes 10 hours, our results suggest that further advances will be required to build such a computer that functions at a practical speed.

The design of the system is shown in Fig. 1(a). The information flow in one cycle is shown in Fig. 1(b), and the full operation is shown schematically in Fig. 1(c).

**Figure 1:**
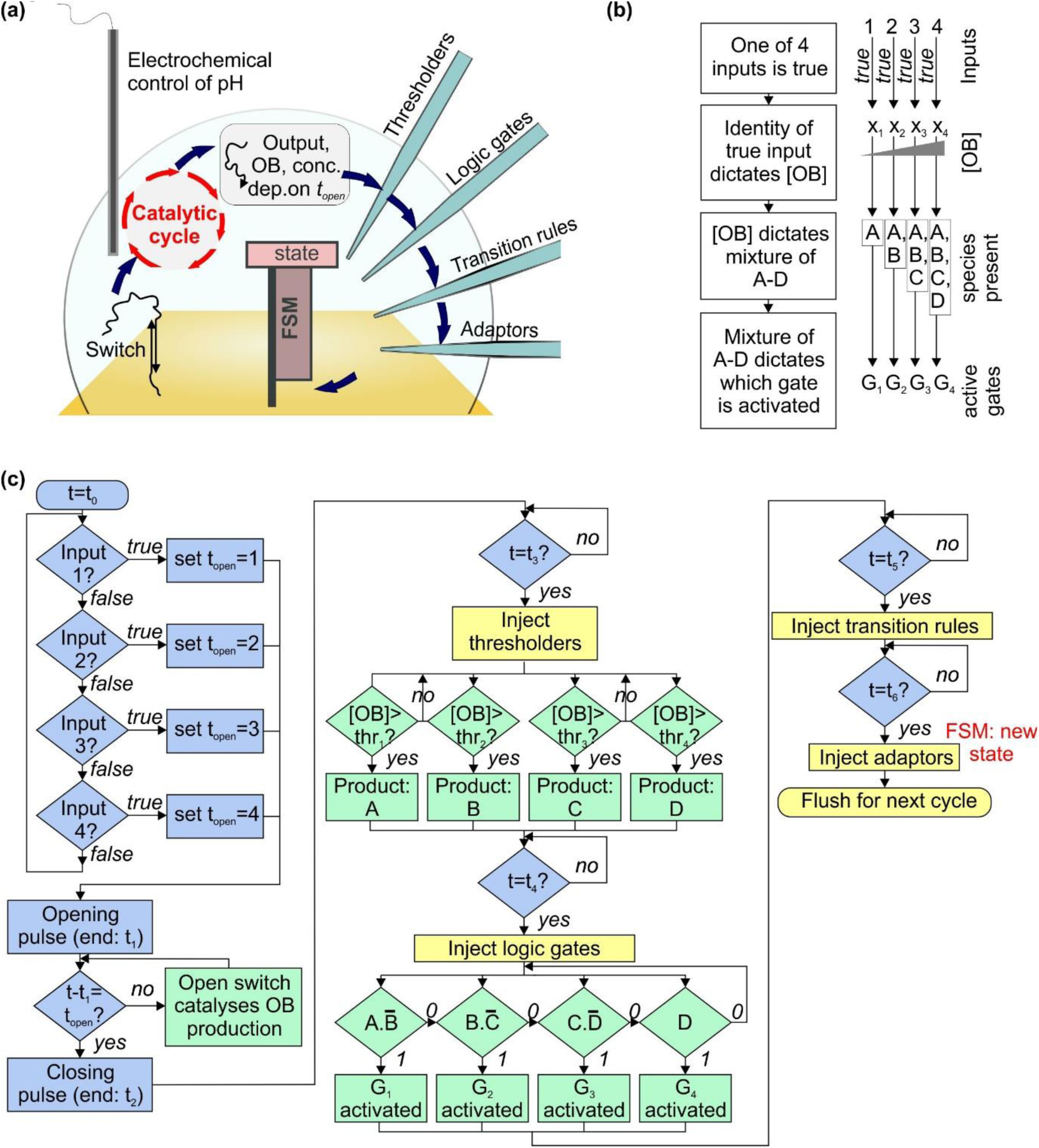
design for a bioelectronic processor that takes electronic inputs and makes decisions autonomously using biomolecular machines. **(a)** Schematic diagram of the system, described fully in the text. The switch is a pH-sensitive DNA nanomachine that changes conformation when the pH is altered electrochemically. Opening the switch reveals a sequence that acts as an catalyst for a cycle converting fuel and substrate molecules into waste product, signal and output. The concentration of the output is assessed using a thresholding mechanism, and the result is then assessed using logic gates. Activated logic gates act as inputs for the finite state machine, and transition rules contain instructions that specify the next state for the machine based on the previous state and the input. **(b)** Flowchart showing how information is encoded in different molecular species within the system. **(c)** Flowchart illustrating the mechanisms of system operation. The boxes are colour-coded by type of process as follows: green – biomolecular, blue – electronic, yellow – microfluidic.

The initial inputs to the system could be the signals from four electronic sensors (or two sensors with dual-rail logic). One and only one of the inputs is true. The identity of the true input is used to set the delay between voltage pulses that are used to electrochemically increase the pH and then return it to its original value.

In our system, the voltage pulses cause conformational changes in pH-sensitive DNA nanoswitches, which are assumed to be based on a triplex^20^ motif and could either float freely in solution or be immobilized on a surface. It has already been shown that the pH at which triplex switches undergo the triplex-duplex transition can be tuned by the choice of sequence employed^21^, which means that our design supports a range of operating conditions.

As shown in Fig. 1(a), open DNA switches catalyse the operation of a cycle^22^ that uses toehold-mediated strand displacement^23,24^ reactions to convert DNA ‘substrate’ and ‘fuel’ molecules into ‘output’ (OB) and ‘signal’ molecules. Assuming there is only one species of switch present, and one set of catalytic molecules, it is not possible to assign one output to each input. Instead, the identity of the true input determines the quantity produced of the output OB, through the delay between opening and closing voltage pulses. The longer this delay, the longer the switch is open, and the more times the catalytic cycle occurs, leading to a higher concentration of OB.

The concentration of OB then determines what mixture of species A-D is left after the thresholder molecules have been injected and allowed to react. The combination of thresholder outputs A-D is assessed using injected logic gates G_1-4_. A-D react with the gates, activating or deactivating them according to the gate design. The parameters are set such that only one gate is active in strength at the end of the process.

Activation of a gate enables it to act as the input for an ensemble of finite state machines immobilized on a surface. Injected transition rule molecules then determine the final state of the machine^9,10^, based on the initial state and the input provided by the logic gates.

Throughout this process, timing of events (i.e. the ‘clock’ of the system) is provided by the underlying electronics, which control the start time and duration of the initial voltage pulse, in addition to the timing of microfluidic injection of thresholders, logic gates, transition rules and adaptors. The latter molecules prepare the finite state machines for the next round of the loop.

## RESULTS

### A catalytic cycle with an electrochemically controlled pH-sensitive nanoswitch as the catalyst

The application of a voltage to a sufficiently concentrated aqueous salt solution leads to electrolysis of water, with pH decreasing at the anode and increasing at the cathode. This effect is reversible, and can be used to modify the pH around an electrode by several units if desired. Other electrochemical reactions can be used in a similar way and it has already been shown that it is possible to control the conformation of pH-sensitive DNA structures electrochemically^14,15^. Depending on the pH of the solution, the nanoswitch is either open or closed. When the switch is closed, a short sequence is sequestered in the loop. This sequence is inaccessible; it is not able to interact with its complements until the switch is opened. The principle of concealing information in this way is extremely well-established^25^.

When the switch is opened, the loop sequence can act as a toehold for downstream strand displacement reactions. However, for many applications it is undesirable to permanently inactivate switches by converting them into waste products, so it is necessary to employ a catalytic cycle in which the nanoswitch catalyst is regenerated. Various approaches could be employed, but the archetypal example of a catalytic cycle is that demonstrated by Zhang *et al.*^22^. This cycle has been characterized thoroughly and reliable models exist for the kinetics of all reactions in solution. Consequently, this is the cycle we choose for our proposed system, and we replace the original linear catalyst strand with the pH-sensitive nanoswitch (Fig. 2(a)). We assume that the nanoswitch is based on a design of Idili *et al.*^21^, and undergoes a transition at pH 7.5. We use the solution-phase kinetic parameters.

**Figure 2:**
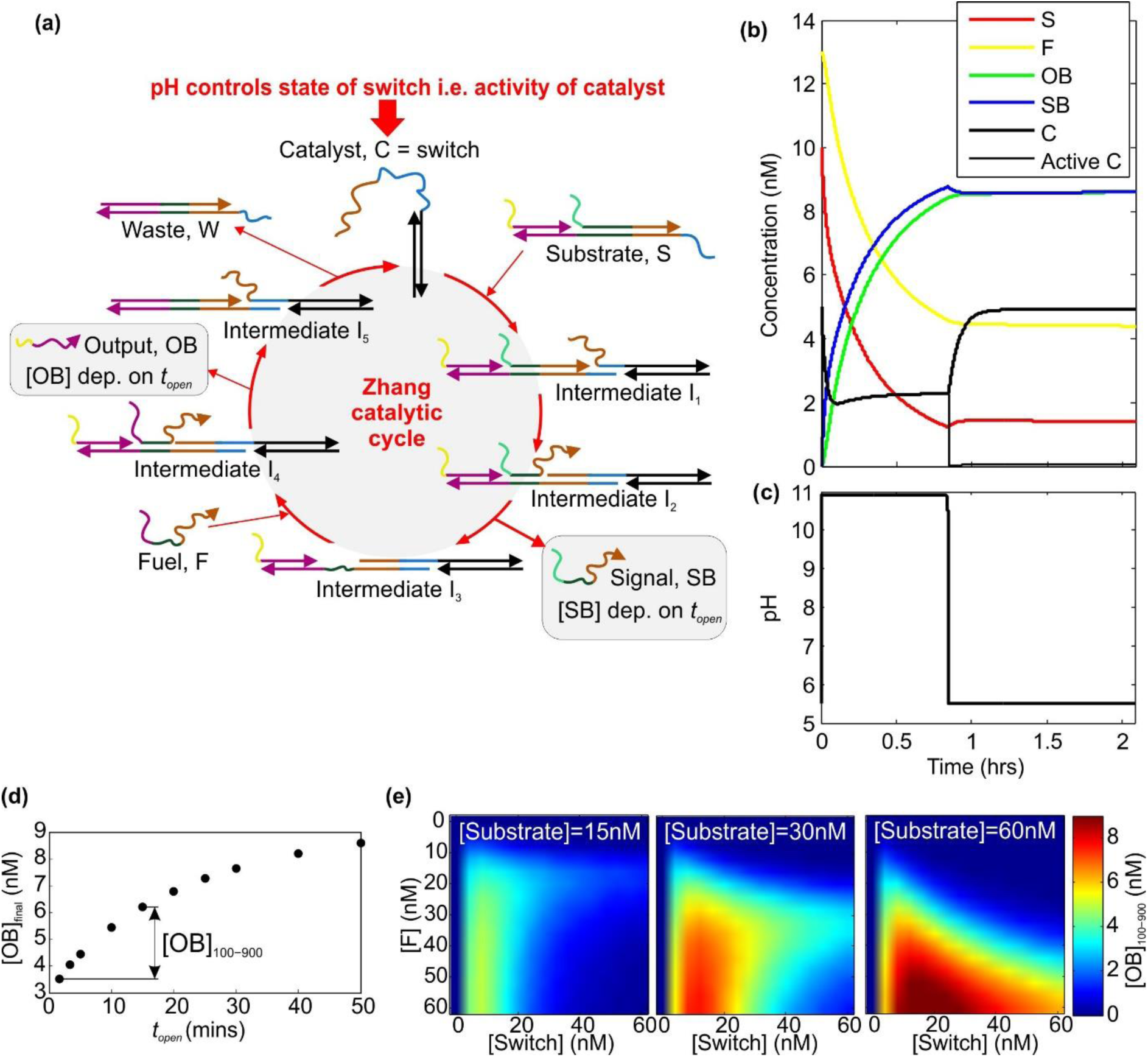
the Zhang catalytic cycle with a pH-sensitive nanoswitch as the catalyst. **(a)** Schematic diagram of the Zhang cycle, based on Fig. 1B from Ref. 22 (adapted with permission from AAAS), with the linear catalyst replaced by a nanoswitch based on the design of Idili *et al*^21^. **(b)** Simulated dynamics, computed using the reaction rates and information provided by Zhang *et al.*^22^. Here, the initial concentrations are: substrate – 10nM, fuel – 13nM, switch – 5nM. For the case illustrated, the separation of the voltage pulses is 3000 seconds, which means that the switch remains open for approximately 50 minutes. The voltage pulses lasted for 20 seconds, and had an amplitude of 4V. The reaction volume was 100μL. **(c)** The pH dynamics corresponding to the simulation in part (b). **(d)** The range of [OB]_final_ values accessible for particular values of t_open_, where t_open_ is assumed to be equal to the temporal separation of opening and closing voltage pulses. The concentrations and voltage pulse characteristics here are the same as for part (b) and (c). **(e)** Illustration that the parameters are inter-dependent. [OB]_100-900_ is defined as shown in part (d), and represents the difference in [OB]_final_ values obtained for *t*_*open*_=100 seconds and *t*_*open*_=900 seconds. A large [OB]_100-900_ value implies a steep curve in the plot shown in (d).

Effectively, this system implements a ‘while’ loop. While the pH is high, SB and OB are produced. The simulated dynamics of this system are shown in Fig. 2(b), and the corresponding pH is shown in Fig. 2(c). The simulation essentially involves the solution of a set of coupled differential equations, as described by Zhang *et al.*, with the addition of an activity variable that defines whether the switch is active as a catalyst (i.e. open) or inactive (i.e. closed). This variable depends on the pH, which is defined by voltage pulses. It will be seen that after the switch has been opened the catalytic cycle can begin operating, with the result that fuel and substrate concentrations fall. The concentration of output OB and signal SB rise, and these curves are very similar. If there is plenty of fuel and all reactions run to completion, the reactions produce one OB for every SB. When the pH is reset, the activity of the catalyst falls to almost zero and the production of OB and SB slows or stops. The concentration of closed switch is restored to its original value. (Note: throughout this paper, the concentration of a species, ‘X’, is denoted using square brackets e.g. [X].)

The final concentration of the output, [OB]_*final*_, is shown in Fig. 2(d) as a function of the period for which the switch is open, for one set of parameters. As *t*_*open*_ increases, [OB]_*final*_ increases, but at a decreasing rate. The range of [OB]_final_ values obtained for a given range of opening times can be changed by altering the concentration of switch, fuel and substrate, as shown in Fig. 2(e). Here, the quantity [OB]_100-900_ is the difference in the final [OB] values (125 minutes after the start of the initial voltage pulse) for t_open_ = 100 seconds and t_open_ = 900 seconds. The data reveals that for a given concentration of substrate and fuel, there is a particular concentration of switch catalyst for which [OB]_100-900_ is maximal. Increasing the switch concentration beyond this value leads to a decrease in [OB]_100-900_. For higher concentrations of substrate and fuel, the final concentration of OB is higher.

The simulation shows that the parameters are interlinked – if one is changed then others must also be changed to ensure optimal performance, where the optimum is defined according to the application. In this paper, we assume an operating range of OB concentrations of 1.5-5nM, which can be achieved using the following parameters: switch catalyst concentration of 1.7nM, substrate and fuel concentrations of 11nM, t_open_ ranging from approximately 1 minute to approximately 40 minutes, total reaction time 125 minutes.

### Using a thresholding mechanism to assess the concentration of the output from the catalytic cycle

The thresholding system is shown in Fig. 3(a), and resembles some of the thresholding mechanisms that have been reported previously^2,5^. Here, the complexes A.L, B.L, C.L and D.L are present in considerable excess. The output OB at concentration [OB]_final_ can displace the species A-D from the strand L through toehold mediated strand displacement, releasing [OB]_final_ /4 of each of A-D. Each of the strands thr_1_-thr_4_ can then bind specifically to one of A-D, rendering it inactive. Thus, in order for one of the species A-D to be observed in solution, [OB]_final_ must exceed 4x the concentration of the corresponding thr. In our simulations, we set the concentrations of thr_1_-thr_4_ to be 0.25, 0.5, 0.75 and 1nM respectively. In this situation, if we wish to produce free A-C, but no free D, we need [OB]_final_ to lie below 4[thr_4_]=4nM but above 4[thr_3_]=3nM. The higher [OB]_final_ lies within this range, the larger the signal from A-C. Fig. 3(b) shows how the concentration of species A-D changes as a function of time for four values of [OB]_final_, with the parameters given here.

**Figure 3:**
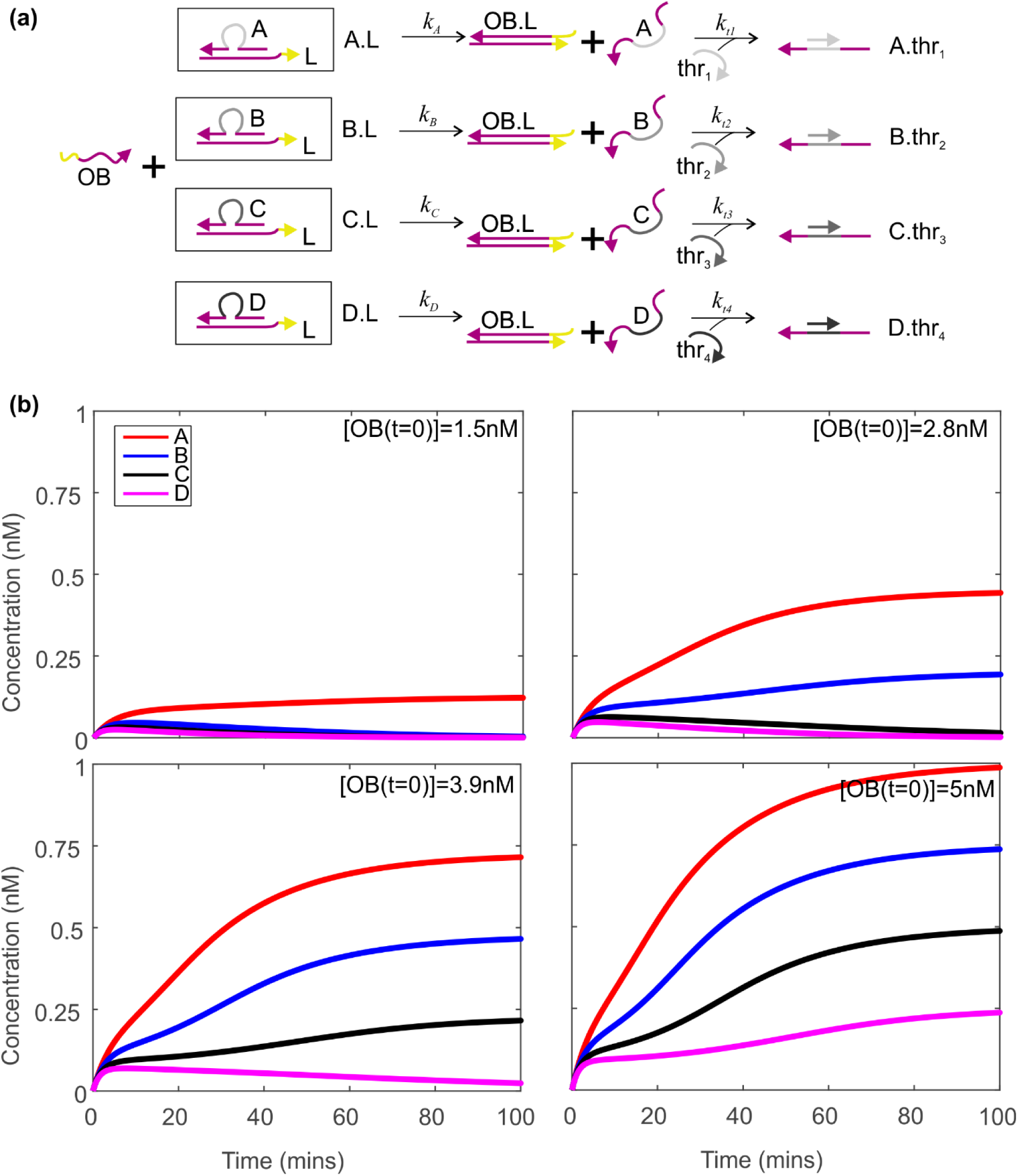
using a thresholding mechanism to assess how much output has been produced by the Zhang cycle. **(a)** The thresholding mechanism. Zhang cycle output OB displaces A-D from the initial complexes. Single-stranded molecules thr_1_-thr_4_ bind to the central (grey) domain of A-D. If the concentration of a thr strand exceeds the concentration of its target A-D, the target species will be entirely consumed by thr. If the target species concentration is greater than that of its corresponding thr, some of the target will be present free and unbound. **(b)** Dynamics of A-D as a function of time for four different concentrations of [OB]. Here, the initial concentrations of thr_1-4_ are 0.25, 0.5, 0.75, 1nM respectively. The total concentrations of A-D are 20nM. Rate constants k_A_-k_D_ are all equal to 1×10^4^ M^−1^s^−1^, and k_t1_-k_t4_ are equal to 1×10^7^ M^−1^s^−1^. 2×10^5^ time steps were used, with a spacing of 0.05 seconds.

### Activation and deactivation of dumb-bell logic gates

Within the system we have defined, there are four scenarios we need to consider: (i) A-only, (ii) A-B with no C and no D, (iii) A-C and no D, (iv) A-D all present. These situations can be distinguished by simple logic operations. The first may be identified by the case A AND NOT B = TRUE. For case ii, the characteristic condition is B AND NOT C = TRUE. The third case corresponds to C AND NOT D = TRUE, while the fourth case is simply D = TRUE. (Each condition is initially characterized by a four-variable statement, but these can be reduced to statements of two variables when it is noted that the four scenarios listed are the cases to be considered. For instance, in this system, it is not possible to have A and C with no B.)

To distinguish these four situations, we propose the use of dumb-bell hairpin logic gates – unimolecular constructs that are activated by one input and deactivated by the second (as shown in Fig. 4(a) for inputs A and B). Static dumb-bell hairpins were used by Rothemund as a marker on the surface of DNA origami tiles^26^. The loop of each hairpin conceals three domains. The finite state machine binding domain is the same for all gates and allows the gate to bind to finite state machines as input. The gate identity domain is the sequence that specifies what the input is, and is unique for each gate. The lock domain may or may not be the same for all gates. On one side of the dumb-bell the sequences in the hairpin loop are the reverse complements of those on the other side.

**Figure 4.**
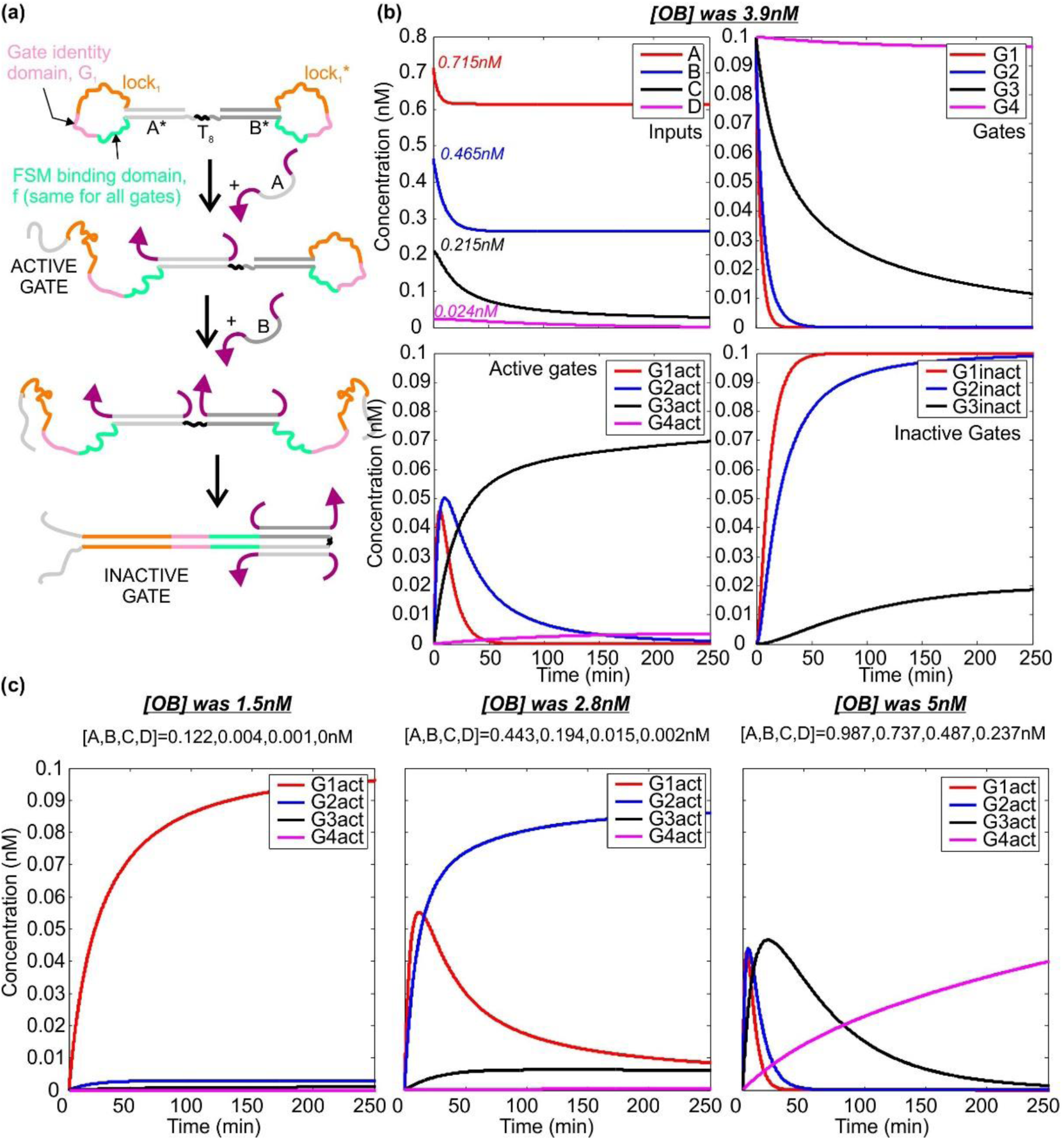
Dumb-bell logic gates – activation and deactivation. (**a**) Operating mechanism of the gates. Binding of species A activates the gate by revealing its identity and lock domains. Binding of species B reveals the reverse complement of both of these domains, leading to gate deactivation. This can occur through the mechanism of toehold-mediated strand displacement even if the FSM-binding domain has already bound to its target. Gates that respond to other pairs of species differ in the sequence of the domains. (**b**) Simulated dynamics for the molecules involved in thresholding, for the case in which the concentration of OB was 3.9nM, and the initial concentrations of A-D were 0.715, 0.465, 0.215 and 0.024nM respectively. Plots show the behaviour of the inputs, the unreacted gates, the active gates and the inactive gates, as labelled. Gate 1 is activated by A and deactivated by B. Gate 2 is activated by B and deactivated by C. Gate 3 is activated by C and deactivated by D. Gate 4 is activated by D and is not deactivated. Gates are present at 0.1nM. The activation and deactivation rates for Gates 1 and 2 are all 6×10^6^ M^−1^s^−1^, the maximum for toehold-mediated strand displacement. For Gate 3, the corresponding rates are 2.5×10^6^ M^−1^s^−1^, and for Gate 4 the activation rate is 2.5×10^5^ M^−1^s^−1^. **(c)** Dynamics of the active gates for three other concentrations of [OB]. In each case, one particular logic gate is activated and the others either do not react or become inactivated.

When the first species binds to the gate, it opens the stem on one side of the dumb-bell. When the second species binds, it opens the other stem. This hairpin loops are complementary and therefore hybridize. It is important to note that this dumb-bell hairpin could not be synthesized as a single strand, because in the absence of any constraints it would assemble incorrectly, with the lock, identity and FSM binding domains bound to each other from the start. However, the dumb-bell could be prepared by straightforward ligation of two separate hairpin constructs.

Fig. 4(b) shows how the species A-D interact with the gates. In this simulation, the initial concentrations of A-D are taken from the thresholder simulation, and all gates are assumed to be present at 0.1nM. The rate at which species A and B displace the corresponding domains in gate 1 is 6×10^6^ M^−1^s^−1^, as is the rate at which B and C displace their targets in gate 2. Species C and D bind to gate 3 at a rate of 2.5×10^6^ M^−1^s^−1^, while D bound to gate 4 more slowly, with a rate constant of 2.5×10^5^M^−1^s^−1^. These rate constants are based on published figures for the rates of toehold mediated strand displacement^23,27^. and hybridization^28^ and can be tuned by modifying factors such as domain length, GC-content and complementarity (i.e. the presence of mismatches).

As the data shows, the concentrations of A-D fall after introduction of the logic gates. The concentration of unreacted gates also falls, and the rate depends on the concentration of the inputs. The dynamics of the active gates depend on the combination of the inputs, as intended. In the example shown in Fig. 2(b), we wanted gate 3 to be activated and the other gates to remain unreacted or be inactivated. It will be seen that this is the result obtained, with very low levels of active gates 1, 2 and 4. Some gate 3 is inactivated but most remains active.

The three other scenarios are shown in Fig. 4(c). In each case, only one gate remains strongly activated, and the concentration of other active gates remains low. This is exactly the desired behaviour. In these simulations, the final concentration of the activated target gate ranges from 0.04 to 0.1nM.

### Operating a finite state machine using the dumb-bell logic gates as input

The result of the steps described so far is the production of a single species – the active gate. For the next step, the transition rules are injected. These are molecules that dictate what the final state of the finite state machines should be, based on the initial state and the input provided by the active gate. One possible molecular implementation of this system is shown in Fig. 5a. The design is based on that of Costa Santini *et al.*^10^, but here the finite state machine is immobilized on the surface and there is no molecular clock. In our system activated dumb-bell hairpins can bind to the immobilized state machines, and the transition rule then binds to the domain formed by the conjunction of the initial state domain of the state machine and the identity domain of the hairpin gate. In principle, at one stage all gates are activated before being inactivated and all active gates can bind to state machines. However, inactivation of the gate will result in its removal from the state machine by strand displacement, and it is not possible for a state transition to occur in the intervening time because the transition rules are not supplied until the logic gates have reached equilibrium with the species A-D.

**Figure 5:**
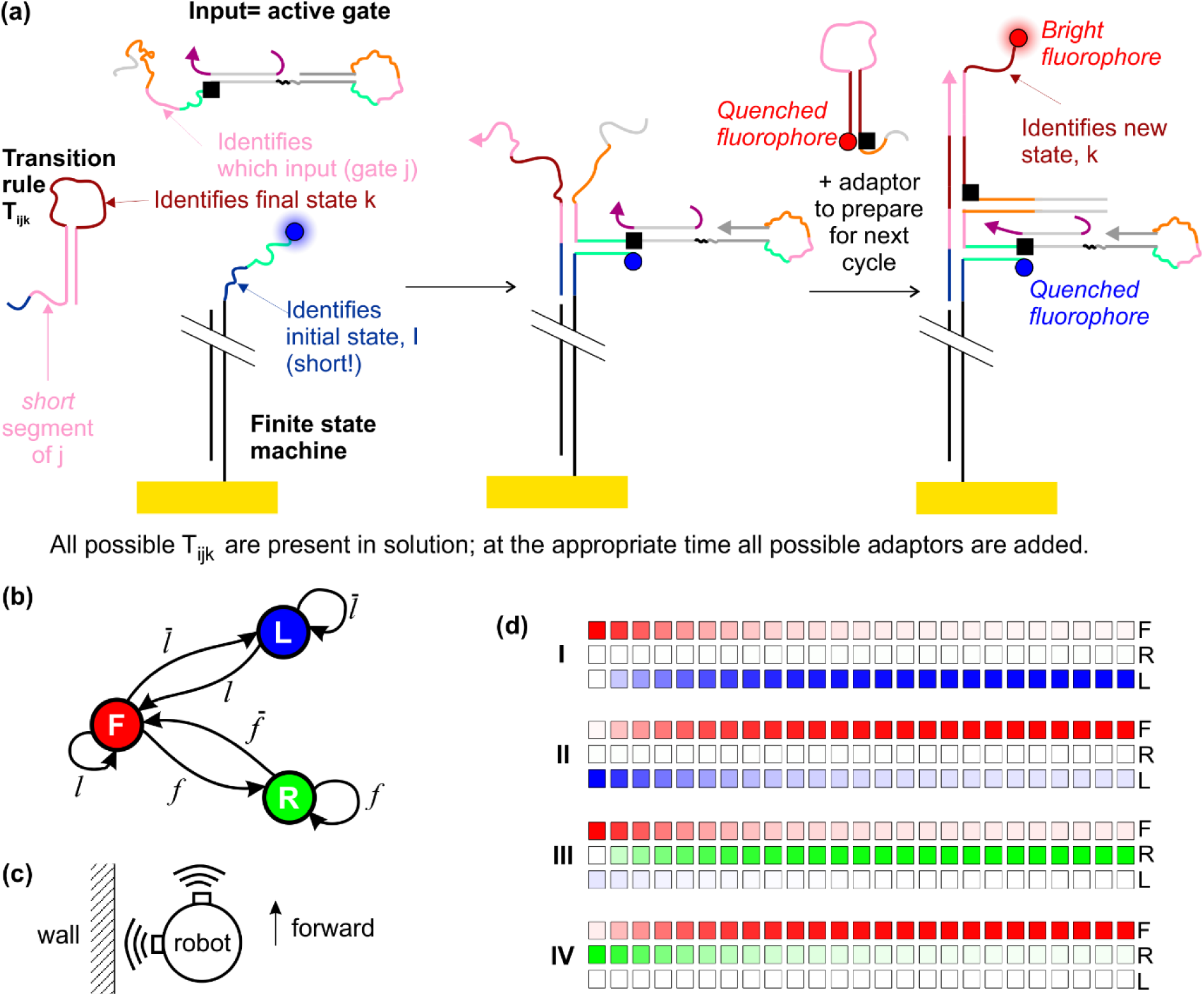
Operating a finite state machine using the dumb-bell logic gates as inputs. **(a)** Molecular implementation of the system, based on the design of Costa Santini *et al*.^10^, with modifications. A possible fluorescence readout scheme is shown. **(b)** Schematic diagram of a finite state machine for a maze-crawling robot that follows the left wall. States are represented in circles and transitions between them are labelled with the condition that induces the transition. **(c)** Schematic diagram of envisaged robot with two sensors. **(d)** Results of simulations modelling four cycles of the system, illustrating how the state changes over time. The results take into account predicted concentrations at the end of every step in each cycle, and are presented in more detail in Table 3. Temporal separation of consecutive squares is 4 minutes.

As a specific example, we may consider the finite state machine that describes the operation of a maze-crawling robot that follows the left wall (Fig. 5(b)). This robot may be in three states – moving forward (F), rotating to the right (R) or rotating to the left (L). These states may be represented by a three-element vector *s=(a,b,c).* In a pure state, a, b and c are either 1 or 0, and only one of them is non-zero. (1,0,0) represents state F, (0,1,0) represents state L and (0,0,1) represents state R. There are two sensors, which register true in the proximity of a wall. One sensor is forward-facing and one faces left. The signal *f* corresponds to a true reading from the forward-facing sensor, and 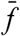 is a false reading from the same sensor. *l* and 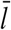 are the corresponding true/false readings for the sensor facing left. We may thus describe the sensors using inputs 1-4 as indicated in Table 1 for representative states and inputs.

**Table 1.** Idealized behaviour of the finite state machine for examples of specified initial conditions and inputs. The state vector contains three elements and is normalized such that it has a magnitude of 1. The input vector is also normalized, but contains four elements, to reflect the four possible input signals.

|  | Initial state | Initial state vector | Signal | True input | Input vector | Final state | Final state vector |
| --- | --- | --- | --- | --- | --- | --- | --- |
| I | F | (1,0,0) | $f$ | 1 | (1,0,0,0) | R | (0,0,1) |
| II | R | (0,0,1) | $\bar{f}$ | 2 | (0,1,0,0) | F | (1,0,0) |
| III | F | (1,0,0) | $\bar{l}$ | 3 | (0,0,1,0) | L | (0,1,0) |
| IV | L | (0,1,0) | $l$ | 4 | (0,0,0,1) | F | (1,0,0) |

The input from the sensors may be represented as a four column vector, *g=(w,x,y,z).* If the input is pure, all but one of the elements is zero and the other element is 1. This also reflects the ideal (normalized) result of the logic gate operation described above. Using index notation and summation convention, the final state *Ψ*_*r*_*=s*_*p*_*g*_*q*_*T*_*pqr*_, where *T*_*pqr*_ is the transition rule. The transition rules are defined in Table 2. *T*_*21r*_, *T*_*22r*_ and *T*_*33r*_ are meaningless and thus denoted by (0,0,0).

**Table 2.**
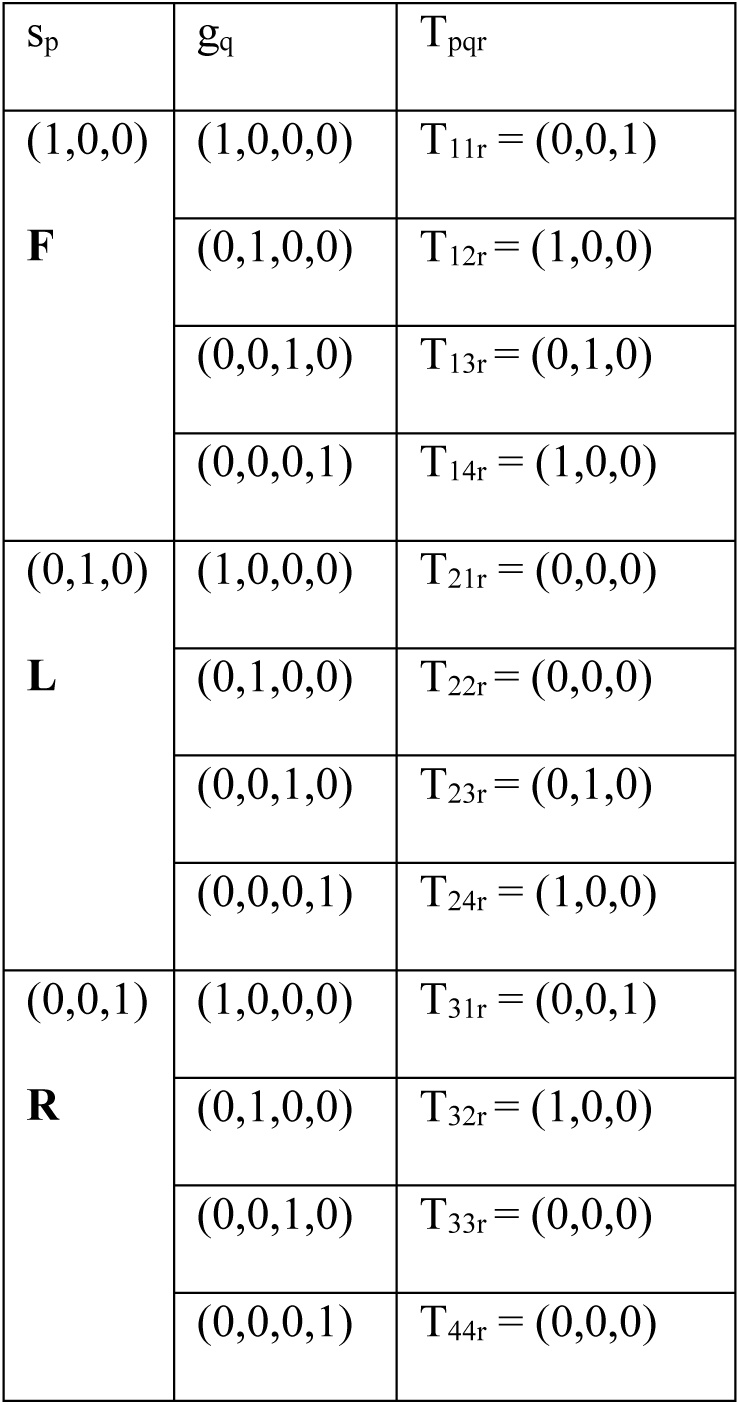
the full set of transition rules for the Finite State Machine illustrated in Figure 5, using the mathematical formalism *Ψ*_*r*_*=s*_*p*_*g*_*q*_*T*_*pqr*_. There are three states and four possible input vectors, which means that 12 transition rules can be mathematically defined. However, four of these rules are irrelevant because the encoded condition could not be observed in reality (T_21r_, T_22r_, T_33r_, T_44r_).

Our simulations can be used to predict the behaviour of the finite state machines for a particular scenario, and the results are shown in Table 3. We assume that the system is initially homogeneous and all machines are in state (1,0,0) i.e. the robot is moving forwards. Next, we assume that the sensors detect a wall in front of the robot. This corresponds to input 1 being true. This is encoded as a specific separation of opening and closing voltage pulses (approximately 1 minute) that gives rise to [OB]=1.5nM. The resulting concentrations of A-D and the active gates are given in the table, together with the computed final state. We consider three subsequent steps, each starting from the final configuration of the previous step. The robot rotates to the right, detects that there is no longer a wall in front of it, moves forward, then detects that there is no longer a wall on the left, rotates to the left, detects that it has found the left wall again and starts moving forward (bottom right entry in table). We assume that the speed of the robot and the range of its sensors are such that action is always taken before the robot hits the wall.

**Table 3.** results of the simulations described in this paper, for the series of states and inputs described in Table 1. The value of [OB] is exact, as is the first initial state. Values of [A]-[D] and [G1]-[G4] are given to 3 d.p. and the state vector elements are given to 4 d.p. Where the state vectors are represented as column vectors, this is for reasons of space rather than mathematical necessity.

|  | Initial<br>state | [OB]<br>(nM) | [A]<br>(nM) | [B]<br>(nM) | [C]<br>(nM) | [D]<br>(nM) | [G<sub>1</sub>]<br>(pM) | [G<sub>2</sub>]<br>(pM) | [G<sub>3</sub>]<br>(pM) | [G<sub>4</sub>]<br>(pM) | Final<br>state |
| --- | --- | --- | --- | --- | --- | --- | --- | --- | --- | --- | --- |
| I | (1,0,0) | 1.5 | 0.122 | 0.004 | 0.001 | 0 | 96.084 | 2.841 | 0.931 | 0 | (0.0296,<br>0.0097,<br>0.9995) |
| II | (0.0296,<br>0.0097,<br>0.9995) | 2.8 | 0.443 | 0.194 | 0.015 | 0.002 | 8.520 | 86.072 | 6.229 | 0.571 | (0.9951,<br>0.0027,<br>0.0985) |
| III | (0.9951,<br>0.0027,<br>0.0985) | 3.9 | 0.715 | 0.465 | 0.215 | 0.024 | 0 | 1.044 | 69.702 | 3.560 | (0.0673,<br>0.9977,<br>0) |
| IV | (0.0673,<br>0.9977,<br>0) | 5.0 | 0.987 | 0.737 | 0.487 | 0.237 | 0 | 0 | 1.377 | 39.846 | (0.9994,<br>0.0345,<br>0) |

Fig. 5(c) represents the results visually, illustrating the dynamics. Here, red denotes state F, green denotes state R, blue denotes state L. As an approximation, we assume that the rate for the change of state is 0.001 s^−1^, based on extrapolation of our previously published experimental data^29^ to the low concentration/surface density regime. We observe that a very high percentage of the finite state machines (over 99%) enter the desired final state.

Although we have considered a specific scenario here, in principle the system we have designed would allow the robot to navigate autonomously through an arbitrary maze without human intervention. For each operating cycle, it would sense its environment using proximity sensors (for example based on infrared radiation) and use a look-up table to automatically encode the sensory input as a particular time delay between opening and closing pulses. Sequential injection of thresholders, logic gates, transition rules and adaptors would follow, where each species would be added automatically at a precise time based on an underlying electronic clock. If the external situation did not change for a long time, a number of operational cycles could pass without a change of state of the finite state machine.

### Readout schemes

For implementation a method of probing the finite state machines will be required. For example, fluorescence signals could be used to indicate the final state. In our framework each adaptor takes the form of a hairpin, which opens upon binding to the finite state machine. Each adaptor could be labelled with a specific fluorophore (e.g. from the Atto family) and an appropriate quencher (e.g. Iowa Black), as illustrated in Fig. 5(a). Opening of the adaptor hairpin would dramatically increase the separation of fluorophore and quencher, leading to a significant rise in the fluorescence signal. The use of multiple colours of fluorophore (one for each state) would allow the output to be read. Of course, it would be necessary to quench the bound fluorophore at the start of the subsequent cycle, to avoid an increase in the background signal. Hence, the input (gate) would also need to be functionalized with a quencher (Fig. 5(a)). To measure the fluorescence, the robot would need an appropriate light source, filters and detectors.

Alternatively an electrochemical strategy could be employed, based on an E-DNA reporter circuit making use of charge transfer between an electrode surface and a redox-active molecule such as methylene blue.

## DISCUSSION

We have presented a possible design for a hybrid bioelectronic computer that processes electrical signals using DNA molecular machines and makes decisions autonomously. Our system has not yet been validated experimentally, but we have performed extensive simulations of its behaviour. Our results suggest that it will operate as expected, and lay the foundations for the construction and testing of a real system. Our own experimental work in this area is ongoing, but we discuss some experimental considerations in Supplementary Discussion 1.

We note that the finite state machines presented in Fig. 5 contain in their structure a memory of their entire history. In principle an add-on module could be designed that would deconstruct the machines and read back the operations performed, allowing the system to repeat its actions in reverse.

In this work, we considered the possibility of using our system to control a maze-crawling robot. This would be a proof-of-concept demonstrator, and the principles could subsequently be expanded to other situations. Our approach is likely to be of particular utility in situations where both biomolecular and electronic stimuli are required. However, as presently envisaged, our system would take 10 hours for each step of the finite state machine, due to the speed of the DNA reactions (Supplementary Discussion 2). This may be prohibitive for many applications, and it would be highly desirable to increase the reaction rate by three or four orders of magnitude – reducing a ten-hour process to a few seconds.

Slight increases in speed could potentially be achieved by optimization of the operating parameters^23,24,27^ such as the concentration of various species and design of DNA sequences. However, increasing the processing speed to the desired levels would require a paradigm shift. For example, Qian and Winfree^5^ have pointed out that the phenol emulsion reassociation technique^30^ may be a viable route for increasing the speed of DNA computers, perhaps by up to four orders of magnitude. Organic solvents can also increase the speed of hybridization^31^. Alternatively, it may be possible to use methods developed to enhance the speed of biochemical assays involving DNA, such as isotachophoresis^32,33^. Furthermore, it has also been shown that recombination proteins may be able to significantly enhance the rate of hybridization^34,35^, and such proteins may have the potential to speed up DNA computers.

For future development of hybrid bioelectronic computers that contain large numbers of interacting DNA strands it will also be necessary to optimize the sequences to minimize spurious interactions and leak reactions. It may be possible to do this by using evolutionary algorithms to identify the most appropriate sequences^36^. Alternatively, DNA circuits could be localized on a surface such as a DNA origami tile, which would allow re-use of sequences, minimizing leak reactions^37^. This approach could also increase reaction speed in comparison with systems that have freely diffusing components.

In DNA computing, many systems are characterized by ‘one-pot’ reactions, where all reactants are present at the same time in the same volume. Our approach does not follow this paradigm. In our proposed system, multiple steps must occur within each information processing layer before the final product is passed to the next layer. If we attempted to employ a one-pot procedure, intermediates from one layer could react prematurely with components from the next layer. For instance, in a one-pot scenario the species ‘A’ produced by the top thresholding reaction in Fig. 3(a) could either interact with the species thr, or the dumb-bell hairpin logic gate shown in Fig. 4(a). In this case, the thresholder circuit would not function correctly.

In principle, the effects of cross-reactivity could be reduced by careful selection of domain lengths and sequences, but eliminating the requirement to use a one-pot procedure allows more flexible design of DNA molecules and opens up the design space for multi-layer information processing systems, enabling a more modular approach.

Our system is limited in terms of speed and takes 10 hours to complete a cycle. One-pot approaches might be an order of magnitude faster, but one hour would still be too slow for many applications. Thus, a ‘one pot’ procedure may have a speed advantage, perhaps by an order of magnitude, but is still too slow for many applications. This means that significant advances will be needed to speed up DNA computers for practical use even if they do employ a one-pot approach.

Furthermore, we note that fundamental information processing operations in biology frequently do not take place in a ‘one-pot’ reaction. For example, we may consider gene expression in eukaryotes. In these organisms, transcription occurs in the nucleus and mRNA transcripts must exit through nuclear pores so that translation can occur at the ribosomes. This multi-layer processing mechanism allows eukaryotes to utilize more complex mechanisms for gene regulation than their prokaryotic counterparts. Taking inspiration from Biology, we suggest that departure from the one-pot reaction paradigm will provide new opportunities for DNA computing, with advantages that outweigh the limitations.

If the speed of the molecular processes can be increased significantly and steps taken to minimize leak reactions, DNA-based bioelectronic computers will have considerable potential. In principle, coupling electronic circuitry and biomolecular devices could enable the construction of low-power biocompatible thinking machines that process information in a highly parallel and potentially fault-tolerant manner. Here we have presented theoretical testing of an initial design for a basic hybrid system, illustrating how information could be transmitted seamlessly across a bioelectronic interface and processed dynamically for autonomous decision making. Future research will build on this study, and could lead to practical devices that could be used in real-world environments.

## METHODS

All simulations were performed using MATLAB R2013a. Each phase of the cycle was modelled as a separate operation. All code is available in the Supplementary Information (Supplementary Methods).

### Calculating the effect of a voltage pulse on the pH-sensitive nanoswitch

We assume that the pH changes as a result of the electrolysis of water. We also assume that the current, *I*, is 0.4mA (voltage of 4V, resistance of 10kΩ) and the volume of solution is 100μL. We neglect diffusion time for OH^−^and H^+^ ions, effectively assuming that they are produced uniformly throughout the solution. The number of ions produced in a time interval *dt* is then *n*=*I* d*t* /(*N*_*A*_*e*), where *N*_*A*_ is Avagadro’s number and *e* is the electron charge. The concentration change is *Δc=n/v*, where *v* is the reaction volume. This allows the pH to be calculated. The initial pH is assumed to be 5.5.

The effective p*K*_a_ of the switch under consideration is taken to be 7.5. In accordance with the work of Idili *et al*.^21^, this can be adjusted by changing the switch design. p*K*_a_ is defined as -log_10_(*K*_a_), where *K*_a_=[A^−^][H^+^]/[AH]. Identifying HA as the protonated (closed) form of the switch and A^−^as the deprotonated (open) form, and using the definition of pH, we obtain [Open]/[Closed]=10^pH-p*K*a^. Defining [Total]=[Open]+[Closed] yields the result that [Open]=[Total]/(1+10^p*K*a-pH^). The quantity [Total] is conserved and defined at the start of the experiment.

### Simulating the behaviour of a Zhang catalytic cycle where the catalyst is a pH-sensitive nanoswitch

The voltage is switched on at the start of the simulation, and left on for 20 seconds. After a delay, a voltage of equal and opposite amplitude is applied for the same length of time. Using the result from above, the activity of the catalyst is defined as [Open]/[Total]. This is computed for all time points in advance. The coupled differential equations provided by Zhang *et al.*^22^ are then solved, assuming that only the active catalyst molecules participate in the reaction. With this approach, we assume that a nanoswitch can only reclose when its loop domain is not bound to another molecule. The time interval used is 0.05 seconds and the number of time steps is 150000, corresponding to a total simulation time of 125 minutes. Substrate, fuel and total switch concentrations are varied.

### Modelling the thresholders

The system of thresholders is modelled as follows. To facilitate compact mathematical formulation, OB is represented as x and A-D are referred to as A_m_, where A_1_corresponds to A while A_2_ corresponds to B, etc. Y.Z represents the construct formed by hybridization of Y and Z, where Y and Z may be A_m_, L_m_, th_m_.

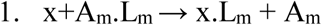

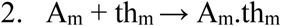

The rate of the first reaction is *k*_*m*_ and the rate of the second is *k*_*tm*_. Here, *k*_*tm*_>>*k*_*m*_. The reactions are irreversible.

The rate of change of free A_m_ is given by:

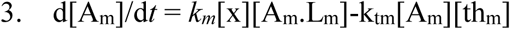

However, we know that the total amount of A_m_ (free or bound to another molecule) is conserved:

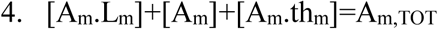

Similarly:

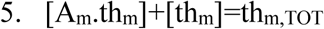

This means that:

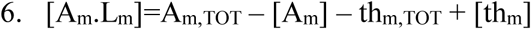

Hence:

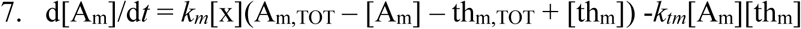

and

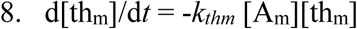

The quantity of output is also conserved, as is the quantity of each L_m_:

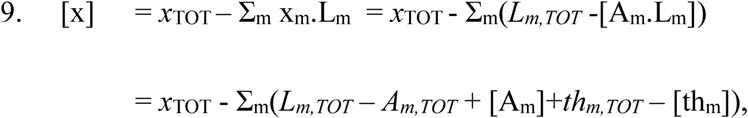

where *x*_TOT_ and *L*_*m,TOT*_ are the conserved quantities.

Using the notation that ℒ= Σ_m_ *L*_*m,TOT*_, 𝒯= = Σ_m_ *th*_*m,TOT*_ and 𝒜 = = Σ_m_ *A*_*m,TOT*_, we obtain:

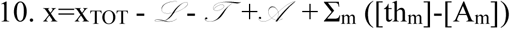

In the simulation, equations 7, 8 and 10 are solved for every time point to yield the dynamics of the species of interest. We assume that for all *m, k*_*m*_=10^4^ M^−1^s^−1^ and *k*_*tm*_=10^7^ M^−1^s^−1^, in accordance with published data^28^. The total concentration of each A_m_ is taken to be 20nM, and the total concentrations of th_1-4_ are 0.25nM, 0.5nM, 0.75nM and 1nM respectively. These parameters could be adjusted as appropriate.

### Simulating the dynamics of the dumb-bell logic gates

The following reactions occur:

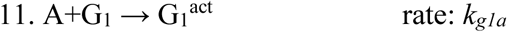

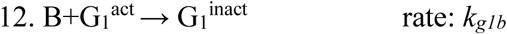

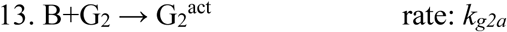

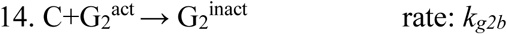

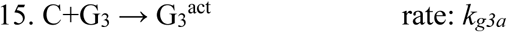

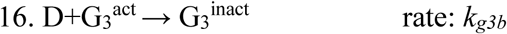

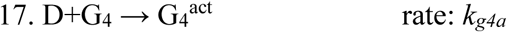

Here, the superscript ‘act’ denotes an activated gate and ‘inact’ denotes an inactivated gate. These reactions give rise to the following set of coupled differential equations, which are solved numerically in MATLAB.

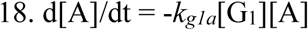

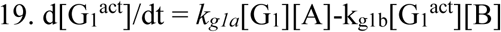

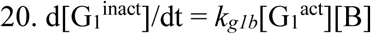

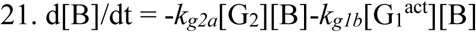

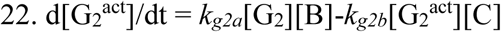

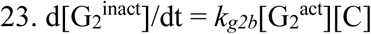

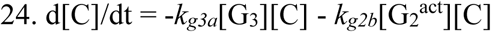

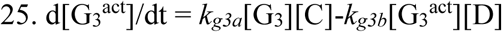

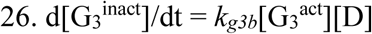

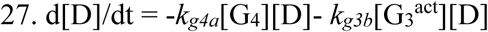

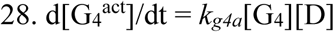

The rate constants are as follows:

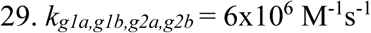

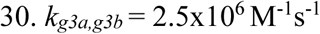

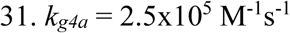

### Simulating the behaviour of the finite state machine

Using index notation and summation convention, the final state of the machine is defined as *Ψ*_*r*_*=s*_*p*_*g*_*q*_*T*_*pqr*,_ where *s*_*p*_ is the initial state, *g*_*q*_ is the input (the activated gate) and *T*_*pqr*_ is the transition rule. Here the state and input vectors are normalized such that |***s***.***s***|*=1* and |***g***.***g***|*=1*.

Here it is not necessary to solve any differential equations numerically because the reaction dynamics can be found analytically. We compute the final state and infer the dynamics by assuming single exponential kinetics. The rate is taken to be 1×10^−3^ s^−1^. This value was estimated based on the results we have obtained previously for hybridization of incoming strands to a surface-immobilized target, with extrapolation to a regime in which the solution concentration is low and the surface density is also low.

## Acknowledgements

The authors thank EPSRC for funding under Platform Grant EP/K040820/1.

## Author Contributions

K.E.D. designed the system and performed the simulations. The manuscript was written by K.E.D., with input from M.A.T, S.J. and A.M.T. All authors gave approval to the final version of the manuscript.

## Additional information

Supplementary Information accompanies this paper.

### Data availability

All the simulation code required to reproduce the results is available in the Supplementary Information.

