## Supplementary Materials for "Towards a bioelectronic computer: A theoretical study of a multi-layer biomolecular computing system that can process electronic inputs"

### Supplementary Discussion 1: Experimental considerations

Our proposed system is modular, and has several layers of information-processing capacity. To test our system experimentally, it would be necessary to test each layer individually and in combination with other layers. The complexity of the tests (in terms of the number of layers involved) would gradually increase until all layers were operating together. As far as possible, our system is based on components that have previously been well-characterized by others.

The operation of individual DNA machines or units within the system could be tested using a scheme in which fluorophore emission is modulated by a quencher or fluorophores interact through FRET. It would not be necessary to perform microscopy, as single-molecule level information is not required. Bulk measurements made using a fluorimeter or equivalent apparatus would suffice. Polyacrylamide gel electrophoresis could also be used to characterize some aspects of the DNA circuitry.

To successfully integrate all layers, it would be necessary to identify a medium that could accommodate all processes. The solution selected should have limited buffering capacity, to enable the pH to be changed electrochemically. The concentration of sodium and/or magnesium ions would need to be at the usual level required for DNA hybridization (mM).

**Supplementary Discussion 2:** How long does each cycle take?

The production of OB takes 125 minutes (including the period for which the switch is open and a subsequent equilibration period). The thresholding stage takes 100 minutes and results in production of some combination of A-D. The operation of the logic gates takes 250 minutes and the finite state machine takes 100 minutes to change state. The total length of each cycle is therefore: 125+100+250+100=575 minutes = 9 hours and 35 minutes.

**Supplementary Methods**

This part of the document provides the MATLAB routines used to perform the simulations described in the paper. The standard MATLAB colour code is used. Underlined code can be edited to change the parameters of the system.

**1. Simulating the Zhang catalytic cycle with a voltage-activated pH sensitive switch as a catalyst**

(a) MATLAB routine – for specific individual concentrations

```
%Katherine Dunn
%Modelling the Zhang catalytic cycle, using a catalyst that is
activated or
%deactivated by voltage pulses
%Note: S=substrate, not signal or switch

close all
clear all

%Definition of time
N=150000;
dt=0.05;
t=[0:dt:N*dt];

%Initial concentration of each species
Sinit=11; %substrate
Finit=11; %fuel
OBinit=0; %output
SBinit=0; %signal
Winit=0; %waste
Cinit=1.7; %catalyst (switch)
I3init=0; %intermediate 3
I5init=0; %intermediate 5
scale=1e-9; %i.e. nM (allows all of the concentrations above to
be written conveniently). Change next line to match this one.
concunit='nM'; %unit for labelling y-axis of graphs

%Reaction rates as given by Zhang
k0=23;
k1=6.5e5;
k2=4.2e5;
k_3=k1;
k_1=k2;
k3=4e-3;

%Definitions and initial values
V=zeros(N+1,1); %Voltage applied is zero for all time by
default, until specified otherwise in code below
```

```
t=[0:dt:N*dt]; %Definition of time

%Definitions
tVon=0; %When is the voltage switched on?
DurationV=20; %How long is the voltage on for?
tVoff=tVon+DurationV; %When is the voltage switched off?
GapV=[60 200 300 600 900 1200 1500 1800 2400 3000 3600]; %How
long after the end of the set pulse does the reset pulse start?
KV=length(GapV);
tVreseton=zeros(KV,1);
tVresetoff=zeros(length(GapV),1);
Von=4; %What is the size of the voltage applied?
NA=6.022e23; %Avagadro's number
e=1.602e-19; %Charge of electron (=size of charge on singly
ionized particle)
R=1e4; %estimate of resistance
volume=100e-6; %Reaction volume. IN LITRES
pHi=5.5; %Initial pH
pH=pHi*ones(N+1,1); %By default, the pH does not change until
the calculation says otherwise
dpH=zeros(N+1,1); %The change in pH is zero until the
calculation says otherwise.
concOHchange=zeros(N+1,1); %The change in the concentration of
hydroxyl ions is zero until calculated.
k_open=ones(N+1,1); %Rate of opening (initialization - values to
be replaced by calculation)
k_close=ones(N+1,1);%Rate of closing (initialization - values to
be replaced by calculation)
Sw_tot=Cinit*scale; %Total concentration of switch
Open_0=0; %Initial concentration of open switch
Open=Open_0*ones(N+1,1); %Initializing variable - concentration
of open switch
pKa=7.5; %pKa of the switch under consideration
Zsig=zeros(1,KV);
Zout=zeros(1,KV);

%Creation of figures
%Figures should occupy full screen
Zfig=figure('units','normalized','outerposition',[0 0 1 1]);
ZOUTfig=figure('units','normalized','outerposition',[0 0 1 1]);
pHfig=figure('units','normalized','outerposition',[0 0 1 1]);
lwi=1.5; %Line width

for kappa=1:KV
    V=zeros(N+1,1);
    tVreseton(kappa)=tVoff+GapV(kappa); %When does the reset
pulse start?
```

```
    tVresetoff(kappa)=tVreseton(kappa)+DurationV; %When does the
reset pulse stop?
    for w=2:N+1
        if t(w)>tVon && t(w)<tVoff
            V(w)=Von; %If we are applying a set pulse at time
point w, let that be reflected in the value of V(w)
        end
        if t(w)>tVreseton(kappa) && t(w)<tVresetoff(kappa)
            V(w)=-Von; %If we are applying a reset pulse at time
point w, let that be reflected in the value of V(w)
        end
        concOHchange(w)=V(w)*dt/(NA*R*e*volume); %This is how
much the concentration of OH- changes in the time interval dt,
assuming constant current
        pH(w)=log10(10^(-14+pH(w-1))+concOHchange(w))+14;%The pH
at this time point is the pH at the last time point plus the
observed change in pH now.
        Open(w)=Sw_tot/(1+10^(-pH(w)+pKa));
    end

    active=Open/Sw_tot;

    %Default values of concentrations for all t are set equal to
values at t=0
    S=scale*Sinit*ones(N+1,1);
    F=scale*Finit*ones(N+1,1);
    OB=scale*OBinit*ones(N+1,1);
    SB=scale*SBinit*ones(N+1,1);
    W=scale*Winit*ones(N+1,1);
    C=scale*Cinit*ones(N+1,1);
    I3=scale*I3init*ones(N+1,1);
    I5=scale*I5init*ones(N+1,1);

    %Until otherwise calculated, there is no change in any of
the
    %concentrations
    dS=zeros(N+1,1);
    dF=zeros(N+1,1);
    dOB=zeros(N+1,1);
    dSB=zeros(N+1,1);
    dW=zeros(N+1,1);
    dC=zeros(N+1,1);
    dI3=zeros(N+1,1);
    dI5=zeros(N+1,1);

    for zT=2:N
```

```
        %The following lines encode the differential equations
that
        %represent the reaction, as described by Zhang, with one
        %modification: the activity of the catalyst is defined
by the value
        %of 'active'.
        dS(zT)=-k0*S(zT)*F(zT)-
k1*active(zT)*S(zT)*C(zT)+k_1*I3(zT)*SB(zT);
        dF(zT)=-k0*S(zT)*F(zT)-k2*I3(zT)*F(zT);
        dOB(zT)=k0*S(zT)*F(zT)+k2*I3(zT)*F(zT);
        dSB(zT)=k0*S(zT)*F(zT)+k1*active(zT)*S(zT)*C(zT)-
k_1*I3(zT)*SB(zT);
        dW(zT)=k0*S(zT)*F(zT)+k3*I5(zT)-
k_3*active(zT)*C(zT)*W(zT);
        dC(zT)=-
k1*S(zT)*active(zT)*C(zT)+k_1*I3(zT)*SB(zT)+k3*I5(zT)-
k_3*active(zT)*C(zT)*W(zT);
        dI3(zT)=k1*active(zT)*S(zT)*C(zT)-k_1*I3(zT)*SB(zT)-
k2*I3(zT)*F(zT);
        dI5(zT)=k2*I3(zT)*F(zT)-
k3*I5(zT)+k_3*active(zT)*C(zT)*W(zT);
        S(zT+1)=S(zT)+dt*dS(zT);
        F(zT+1)=F(zT)+dt*dF(zT);
        OB(zT+1)=OB(zT)+dt*dOB(zT);
        SB(zT+1)=SB(zT)+dt*dSB(zT);
        W(zT+1)=W(zT)+dt*dW(zT);
        C(zT+1)=C(zT)+dt*dC(zT);
        I3(zT+1)=I3(zT)+dt*dI3(zT);
        I5(zT+1)=I5(zT)+dt*dI5(zT);
    end


    %Now I plot the results.
    figure(Zfig)
    subplot(2,ceil(KV/2),kappa)
    plot(t/3600,S/scale,'r','LineWidth',lwi);
    %Time is plotted in hours. The concentration is plotted with
    %reference to the scale defined at the start of the code.
    hold on
    plot(t/3600,F/scale,'y','LineWidth',lwi);
    plot(t/3600,OB/scale,'g','LineWidth',lwi);
    plot(t/3600,SB/scale,'b','LineWidth',lwi);
    %plot(t/3600,W/scale,'m','LineWidth',lwi);
    plot(t/3600,C/scale,'k','LineWidth',lwi);
    %plot(t/3600,I3/scale,'c','LineWidth',lwi);
    %plot(t/3600,I5/scale,'c','LineWidth',lwi-0.5);
    plot(t/3600,active.*C./scale,'k','LineWidth',lwi-0.5);
```

```
    xlim([0 max(t/3600)]);
    xlabel('Time (hrs)');
    ylabel(strcat('Concentration (',concunit,')'));
    Zsig(kappa)=SB(N+1);
    Zout(kappa)=OB(N+1);
    if kappa==KV
        %For the last plot, add a legend. Put the legend on the
right hand side.
        %NB colour code is the same for all plots.
        %legend({'S','F','OB','SB','W','C','I3','I5','Active
C'},'Location','east');
        legend({'S','F','OB','SB','C','Active
C'},'Location','east');
    end
    figure(pHfig)
    subplot(2,ceil(KV/2),kappa)
    plot(t/3600,pH,'k','LineWidth',lwi);
    xlim([0 max(t/3600)]);
    xlabel('Time (hrs)');
    ylabel('pH');
end
figure(ZOUTfig)
scatter(GapV/60,Zout/scale,80,'k','filled');
hold on
%scatter(GapV/60,Zsig/scale,80,'r','filled');
xlabel('t_{open} (mins)');
ylabel(strcat('Final concentration of OB (',concunit,')'));
```

(b) MATLAB routine – for a range of parameters.

```
%Katherine Dunn
%Modelling the Zhang catalytic cycle, using a catalyst that is
activated or
%deactivated by voltage pulses
%Note: S=substrate, not signal or switch
%VARIABLE F, C, s

close all
clear all

%Definition of time
N=150000;
dt=0.05;
t=[0:dt:N*dt];

%Initial concentration of each species
Sinitvar=[15 30 60]; %substrate
Finitvar=[0 4 8 12 16 20 24 28 32 36 40 44 48 52 56 60]; %fuel
OBinit=0; %output
SBinit=0; %signal
Winit=0; %waste
Cinitvar=[0 4 8 12 16 20 24 28 32 36 40 44 48 52 56 60];
%catalyst (switch)
I3init=0; %intermediate 3
I5init=0; %intermediate 5
scale=1e-9; %i.e. nM (allows all of the concentrations above to
be written conveniently). Change next line to match this one.
concunit='nM'; %unit for labelling y-axis of graphs

%Reaction rates as given by Zhang
k0=23;
k1=6.5e5;
k2=4.2e5;
k_3=k1;
k_1=k2;
k3=4e-3;

%Definitions and initial values
V=zeros(N+1,1); %Voltage applied is zero for all time by
default, until specified otherwise in code below
t=[0:dt:N*dt]; %Definition of time

%Definitions
tVon=0; %When is the voltage switched on?
DurationV=20; %How long is the voltage on for?
tVoff=tVon+DurationV; %When is the voltage switched off?
```

```
GapV=[100 900]; %How long after the end of the set pulse does
the reset pulse start?
KV=length(GapV);
tVreseton=zeros(KV,1);
tVresetoff=zeros(length(GapV),1);
Von=4; %What is the size of the voltage applied?
NA=6.022e23; %Avagadro's number
e=1.602e-19; %Charge of electron (=size of charge on singly
ionized particle)
R=1e4; %estimate of resistance
volume=100e-6; %Reaction volume. IN LITRES
pHi=5.5; %Initial pH
pH=pHi*ones(N+1,1); %By default, the pH does not change until
the calculation says otherwise
dpH=zeros(N+1,1); %The change in pH is zero until the
calculation says otherwise.
concOHchange=zeros(N+1,1); %The change in the concentration of
hydroxyl ions is zero until calculated.
k_open=ones(N+1,1); %Rate of opening (initialization - values to
be replaced by calculation)
k_close=ones(N+1,1);%Rate of closing (initialization - values to
be replaced by calculation)
Open_0=0; %Initial concentration of open switch
Open=Open_0*ones(N+1,1); %Initializing variable - concentration
of open switch
pKa=7.5; %pKa of the switch under consideration
Zsig=zeros(1,KV);
Zout=zeros(1,KV);

%Creation of figures
%Figures should occupy full screen
OBfig=figure('units','normalized','outerposition',[0 0 1 1]);
OB100to900=zeros(length(Finitvar),length(Cinitvar));

for si=1:length(Sinitvar)
    Sinit=Sinitvar(si);
for ci=1:length(Cinitvar)
    Cinit=Cinitvar(ci);
    Sw_tot=Cinit*scale; %Total concentration of switch
for fi=1:length(Finitvar)
    Finit=Finitvar(fi);
for kappa=1:KV
    V=zeros(N+1,1);
    tVreseton(kappa)=tVoff+GapV(kappa); %When does the reset
pulse start?
    tVresetoff(kappa)=tVreseton(kappa)+DurationV; %When does the
reset pulse stop?
```

```
    for w=2:N+1
        if t(w)>tVon && t(w)<tVoff
            V(w)=Von; %If we are applying a set pulse at time
point w, let that be reflected in the value of V(w)
        end
        if t(w)>tVreseton(kappa) && t(w)<tVresetoff(kappa)
            V(w)=-Von; %If we are applying a reset pulse at time
point w, let that be reflected in the value of V(w)
        end
        concOHchange(w)=V(w)*dt/(NA*R*e*volume); %This is how
much the concentration of OH- changes in the time interval dt,
assuming constant current
        pH(w)=log10(10^(-14+pH(w-1))+concOHchange(w))+14;%The pH
at this time point is the pH at the last time point plus the
observed change in pH now.
        Open(w)=Sw_tot/(1+10^(-pH(w)+pKa));
    end

    active=Open/Sw_tot;

    %Default values of concentrations for all t are set equal to
values at t=0
    S=scale*Sinit*ones(N+1,1);
    F=scale*Finit*ones(N+1,1);
    OB=scale*OBinit*ones(N+1,1);
    SB=scale*SBinit*ones(N+1,1);
    W=scale*Winit*ones(N+1,1);
    C=scale*Cinit*ones(N+1,1);
    I3=scale*I3init*ones(N+1,1);
    I5=scale*I5init*ones(N+1,1);

    %Until otherwise calculated, there is no change in any of
the
    %concentrations
    dS=zeros(N+1,1);
    dF=zeros(N+1,1);
    dOB=zeros(N+1,1);
    dSB=zeros(N+1,1);
    dW=zeros(N+1,1);
    dC=zeros(N+1,1);
    dI3=zeros(N+1,1);
    dI5=zeros(N+1,1);

    for zT=2:N
        %The following lines encode the differential equations
that
        %represent the reaction, as described by Zhang, with one
```

```
        %modification: the activity of the catalyst is defined
by the value
        %of 'active'.
        dS(zT)=-k0*S(zT)*F(zT)-
k1*active(zT)*S(zT)*C(zT)+k_1*I3(zT)*SB(zT);
        dF(zT)=-k0*S(zT)*F(zT)-k2*I3(zT)*F(zT);
        dOB(zT)=k0*S(zT)*F(zT)+k2*I3(zT)*F(zT);
        dSB(zT)=k0*S(zT)*F(zT)+k1*active(zT)*S(zT)*C(zT)-
k_1*I3(zT)*SB(zT);
        dW(zT)=k0*S(zT)*F(zT)+k3*I5(zT)-
k_3*active(zT)*C(zT)*W(zT);
        dC(zT)=-
k1*S(zT)*active(zT)*C(zT)+k_1*I3(zT)*SB(zT)+k3*I5(zT)-
k_3*active(zT)*C(zT)*W(zT);
        dI3(zT)=k1*active(zT)*S(zT)*C(zT)-k_1*I3(zT)*SB(zT)-
k2*I3(zT)*F(zT);
        dI5(zT)=k2*I3(zT)*F(zT)-
k3*I5(zT)+k_3*active(zT)*C(zT)*W(zT);
        S(zT+1)=S(zT)+dt*dS(zT);
        F(zT+1)=F(zT)+dt*dF(zT);
        OB(zT+1)=OB(zT)+dt*dOB(zT);
        SB(zT+1)=SB(zT)+dt*dSB(zT);
        W(zT+1)=W(zT)+dt*dW(zT);
        C(zT+1)=C(zT)+dt*dC(zT);
        I3(zT+1)=I3(zT)+dt*dI3(zT);
        I5(zT+1)=I5(zT)+dt*dI5(zT);
    end
    Zout(kappa)=OB(N+1);
end
OB100to900(fi,ci)=Zout(2)-Zout(1);
end
end
subplot(3,1,si)
imagesc(Cinitvar,Finitvar,OB100to900/scale);
axis square
caxis([0 9]);
co=colorbar;
ylabel(co,'[OB], 100-900');
xlabel('Concentration of switch (nM)');
ylabel('Concentration of fuel (nM)');
title(strcat('[Substrate]=',int2str(Sinit),'nM'));
end
```

**2. Modelling the thresholding reactions**
MATLAB routine.

```
%Katherine Dunn
%Modelling a set of threshold reactions
%Target species x reacts with the complexes made by A-D with 
their
%partners L1-L4. Species th1-th4 react with A-D. Free A-D left 
in solution
%if concentration of A-D released from L1-4 exceeds initial 
concentration of
%corresponding tA-tD.

close all
clear all

scale=1e-9;
concunit='nM';
x0=[1.5 2.2 2.7 2.8 2.9 3.0 3.6 3.7 3.8 3.9 5 6]*scale; %Initial 
concentration of input (=output from Zhang cycle)

%Rate constants for the four thresholding reactions
kA=1e4;
kB=1e4;
kC=1e4;
kD=1e4;
kt1=1e7;
kt2=1e7;
kt3=1e7;
kt4=1e7;
%Total concentrations of thresholder molecules
Atot=20*scale;
Btot=20*scale;
Ctot=20*scale;
Dtot=20*scale;
th1=0.25*scale;
th2=0.5*scale;
th3=0.75*scale;
th4=1*scale;
%Assuming that all the threshold signal molecules are initially 
bound to
%partners L1-4:
L1tot=Atot;
L2tot=Btot;
L3tot=Ctot;
L4tot=Dtot;

%Definition of simulation time etc
```

```
N=200000; %no of iterations
dt=0.05; %time increment
T=N*dt; %length of time
n=100*60/dt;

%Initializing vectors
t=zeros(N+1,1);
A=zeros(N+1,1);
B=zeros(N+1,1);
C=zeros(N+1,1);
D=zeros(N+1,1);
tA=th1.*ones(N+1,1);
tB=th2.*ones(N+1,1);
tC=th3.*ones(N+1,1);
tD=th4.*ones(N+1,1);

thfig1=figure('units','normalized','outerposition',[0 0 1 1]);
thfig2=figure('units','normalized','outerposition',[0 0 1 1]);

fileID=fopen('threshoutput.txt','w');
fprintf(fileID,'Concentration of free A, B, C, D at t=100
minutes\r\n');
fprintf(fileID,'x0\t A\t B\t C\t D\r\n');

for j=1:length(x0)
    %Trying out different concentration of output (note that
here x is the
    %output of the catalytic cycle)
    xtot=x0(j);
    x=xtot*ones(N+1,1);
    %Definition of constant in DE
    K=xtot+Atot+Btot+Ctot+Dtot-L1tot-L2tot-L3tot-L4tot-th1-th2-
th3-th4;
    for i=2:N+1
        %Solving the differential equations
        t(i)=(i-1)*dt;
        tA(i)=tA(i-1)-kt1*A(i-1)*tA(i-1)*dt; %These are the
thresholders that eat up excess A-D
        tB(i)=tB(i-1)-kt2*B(i-1)*tB(i-1)*dt;
        tC(i)=tC(i-1)-kt3*C(i-1)*tC(i-1)*dt;
        tD(i)=tD(i-1)-kt4*D(i-1)*tD(i-1)*dt;
        dA=kA*x(i-1)*(Atot-A(i-1)-th1+tA(i-1))*dt-kt1*A(i-
1)*tA(i-1)*dt;
        dB=kB*x(i-1)*(Btot-B(i-1)-th2+tB(i-1))*dt-kt2*B(i-
1)*tB(i-1)*dt;
        dC=kC*x(i-1)*(Ctot-C(i-1)-th3+tC(i-1))*dt-kt3*C(i-
1)*tC(i-1)*dt;
```

```
        dD=kD*x(i-1)*(Dtot-D(i-1)-th4+tD(i-1))*dt-kt4*D(i-
1)*tD(i-1)*dt;
        A(i)=A(i-1)+dA;
        B(i)=B(i-1)+dB;
        C(i)=C(i-1)+dC;
        D(i)=D(i-1)+dD;
        x(i)=K-A(i)-B(i)-C(i)-D(i)+tA(i)+tB(i)+tC(i)+tD(i);
    end
%Creating plots
figure(thfig1)
subplot(3,4,j)
%plot(t,x,'k');
hold on
plot(t/60,A/scale,'r','LineWidth',2.0);
plot(t/60,B/scale,'b','LineWidth',2.0);
plot(t/60,C/scale,'k','LineWidth',2.0);
plot(t/60,D/scale,'m','LineWidth',2.0);
xlim([0 100]);
%ylim([0 2]);
ch=['[x(t=0)]=' num2str(xtot)];
title(ch);
xlabel('Time (mins)');
ylabel(strcat('Concentration (',concunit,')'));
figure(thfig2)
subplot(3,4,j)
%plot(t,x,'k');
hold on
plot(t/60,tA/scale,'r','LineWidth',2.0);
plot(t/60,tB/scale,'b','LineWidth',2.0);
plot(t/60,tC/scale,'k','LineWidth',2.0);
plot(t/60,tD/scale,'m','LineWidth',2.0);
ch=['[x(t=0)]=' num2str(xtot)]; %create the title
title(ch);
xlabel('Time (mins)');
ylabel(strcat('Concentration (',concunit,')'));

fprintf(fileID,'%2.1f\t %5.3f\t %5.3f\t %5.3f\t
%5.3f\r\n',x0(j)/scale,A(n)/scale,B(n)/scale,C(n)/scale,D(n)/sca
le);
end
fclose(fileID);
```

**3. Modelling the logic gates**
MATLAB routine.

```
%Katherine Dunn
%This script simulates the behaviour of a set of DNA logic
gates.
%The function of the gates is to produce a single output from a
combination
%of inputs. Gate is active if output is true
%Gate G1: A AND NOT B
%Gate G2: B AND NOT C
%Gate G3: C AND NOT D
%Gate G4: D is present

clear all
close all

%Initial concentrations of A-D
A0=0.987e-9;
B0=0.737e-9;
C0=0.487e-9;
D0=0.237e-9;

%Initial concentrations of gates
G10=0.1e-9;
G20=0.1e-9;
G30=0.1e-9;
G40=0.1e-9;

%Rates
kg1a=6e6;
kg1b=6e6;
kg2a=6e6;
kg2b=6e6;
kg3a=2.5e6;
kg3b=2.5e6;
kg4a=2.5e5;

N=200000; %no of iterations
dt=0.1; %time increment
T=N*dt; %length of time

%Initialization of vectors
A=A0*ones(N+1,1);
B=B0*ones(N+1,1);
C=C0*ones(N+1,1);
D=D0*ones(N+1,1);
G1=G10*ones(N+1,1);
```

```
G2=G20*ones(N+1,1);
G3=G30*ones(N+1,1);
G4=G40*ones(N+1,1);
G1act=zeros(N+1,1);
G2act=zeros(N+1,1);
G3act=zeros(N+1,1);
G4act=zeros(N+1,1);
G1inact=zeros(N+1,1);
G2inact=zeros(N+1,1);
G3inact=zeros(N+1,1);

t=zeros(N+1,1);

for i=2:N+1
    %Numerical solution of coupled differential equations (to
avoid
    %excessive bracketing the multiplication by dt is done in
the last
    %step)
    t(i)=(i-1)*dt;
    dA=-kg1a*A(i-1)*G1(i-1);
    dG1=-kg1a*A(i-1)*G1(i-1);
    dG1act=+kg1a*A(i-1)*G1(i-1)-kg1b*B(i-1)*G1act(i-1);
    dG1inact=kg1b*B(i-1)*G1act(i-1);
    dB=-kg1b*B(i-1)*G1act(i-1)-kg2a*G2(i-1)*B(i-1);
    dG2=-kg2a*G2(i-1)*B(i-1);
    dG2act=kg2a*B(i-1)*G2(i-1)-kg2b*C(i-1)*G2act(i-1);
    dG2inact=kg2b*C(i-1)*G2act(i-1);
    dC=-kg2b*C(i-1)*G2act(i-1)-kg3a*G3(i-1)*C(i-1);
    dG3=-kg3a*G3(i-1)*C(i-1);
    dG3act=kg3a*C(i-1)*G3(i-1)-kg3b*D(i-1)*G3act(i-1);
    dG3inact=kg3b*D(i-1)*G3act(i-1);
    dD=-kg3b*D(i-1)*G3act(i-1)-kg4a*G4(i-1)*D(i-1);
    dG4=-kg4a*G4(i-1)*D(i-1);
    dG4act=kg4a*D(i-1)*G4(i-1);
    A(i)=A(i-1)+dA*dt;
    B(i)=B(i-1)+dB*dt;
    C(i)=C(i-1)+dC*dt;
    D(i)=D(i-1)+dD*dt;
    G1(i)=G1(i-1)+dG1*dt;
    G2(i)=G2(i-1)+dG2*dt;
    G3(i)=G3(i-1)+dG3*dt;
    G4(i)=G4(i-1)+dG4*dt;
    G1act(i)=G1act(i-1)+dG1act*dt;
    G2act(i)=G2act(i-1)+dG2act*dt;
    G3act(i)=G3act(i-1)+dG3act*dt;
    G4act(i)=G4act(i-1)+dG4act*dt;
```

```
    G1inact(i)=G1inact(i-1)+dG1inact*dt;
    G2inact(i)=G2inact(i-1)+dG2inact*dt;
    G3inact(i)=G3inact(i-1)+dG3inact*dt;
end
scale=1e-9;
%Plotting figures. The most significant plot is the one in the
lower left
%corner, which shows the concentration of active gates.
myfig=figure('units','normalized','outerposition',[0 0 0.5 1]);
subplot(2,2,1);
plot(t/60,A/scale,'r','LineWidth',2.0);
hold on
plot(t/60,B/scale,'b','LineWidth',2.0);
plot(t/60,C/scale,'k','LineWidth',2.0);
plot(t/60,D/scale,'m','LineWidth',2.0);
title('Inputs')
legend({'A','B','C','D'})
xlabel('Time (min)');
ylabel('Concentration (nM)');
xlim([0 250]);
subplot(2,2,2);
plot(t/60,G1/scale,'r','LineWidth',2.0);
hold on
plot(t/60,G2/scale,'b','LineWidth',2.0);
plot(t/60,G3/scale,'k','LineWidth',2.0);
plot(t/60,G4/scale,'m','LineWidth',2.0);
title('Gates')
legend({'G1','G2','G3','G4'})
xlabel('Time (min)');
xlim([0 250]);
ylabel('Concentration (nM)');
ylim([0 0.1]);
subplot(2,2,3);
plot(t/60,G1act/scale,'r','LineWidth',2.0);
hold on
plot(t/60,G2act/scale,'b','LineWidth',2.0);
plot(t/60,G3act/scale,'k','LineWidth',2.0);
plot(t/60,G4act/scale,'m','LineWidth',2.0);
title('Active gates')
legend({'G1act','G2act','G3act','G4act'})
xlabel('Time (min)');
xlim([0 250]);
ylim([0 0.1]);
ylabel('Concentration (nM)');
subplot(2,2,4);
plot(t/60,G1inact/scale,'r','LineWidth',2.0);
hold on
```

```
plot(t/60,G2inact/scale,'b','LineWidth',2.0);
plot(t/60,G3inact/scale,'k','LineWidth',2.0);
legend({'G1inact','G2inact','G3inact'})
ylim([0 0.1]);
title('Inactive Gates')
legend('show')
xlabel('Time (min)');
xlim([0 250]);
ylabel('Concentration (nM)');

g1a_250=G1act(150001);
g2a_250=G2act(150001);
g3a_250=G3act(150001);
g4a_250=G4act(150001);
```

**4. Modelling the finite state machine**
MATLAB routine.

```
%Katherine Dunn
%Simulation of a DNA finite state machine changing state.
%This code takes one initial state and one condition and shows
how the
%machine will transition to the final state.
%The states and conditions do not have to be 'pure', so this is
really a
%simulation of an ensemble of machines, some of which may be in
the wrong state.
%Each 'pure' state corresponds to a single output colour that
could be aligned
%with fluorescence of e.g. Alexa Fluor 568 (red), 488 (green)
and 350 (blue).

clear all
close all

%Numbers specifying initial state of machine and identifying
which gates
%are active.
s1=0.0673;
s2=0.9977;
s3=0;
g1=0;
g2=0;
g3=1.377;
g4=39.846;

%Definition of a vector specifying the initial state
st_in=[s1 s2 s3]/norm([s1 s2 s3]);

%Definition of a vector specifying which gates are activated
G=[g1 g2 g3 g4]/norm([g1 g2 g3 g4]);

%Definition of time vector. Note that this code does not solve a
%differential equation numerically, and therefore it is not
necessary for
%the time points to be finely spaced.
N=25;
dt=240;
t=0:dt:N*dt;

%The rate at which the state changes. Assume single exponential
kinetics.
kcol=0.001;
```

```
T=zeros(3,4,3);
%The elements of this array specify the transition rules.
%Let t_r = sum over p and q of state_p gate_q M_pqr
%t_r is then a vector that defines the final state.
%The first element of the vector t_r is the amplitude in the
'red' state
%The second element of the vector t_r is the amplitude in the
'green' state
%The third element of the vector t_r is the amplitude in the
'blue' state
T(1,1,:)=[0 0 1];
T(1,2,:)=[1 0 0];
T(1,3,:)=[0 1 0];
T(1,4,:)=[1 0 0];
T(2,3,:)=[0 1 0];
T(2,4,:)=[1 0 0];
T(3,1,:)=[0 0 1];
T(3,2,:)=[1 0 0];
v=ones(3,1);
w=zeros(3,1);


for p=1:3
    for q=1:4
        v(:)=st_in(p)*G(q)*T(p,q,:); %This line calculates the
contribution from each gate, state and transition rule to the
final state
        w(:)=w(:)+v(:); %This is the sum over all contributions
(the code allows impure initial states)
    end
end

w=w/norm(w);

red_i=st_in(1);
green_i=st_in(2);
blue_i=st_in(3);

red_t=ones(1,25);
green_t=ones(1,25);
blue_t=ones(1,25);

red_f=w(1);
green_f=w(2);
blue_f=w(3);
```

```
colours=figure('units','normalized','outerposition',[0 0.2 1
0.3]);

for fcount=1:N
    %This loop calculates the strength of red, green or blue
fluorescence
    %at the timepoint specified by the loop iteration number.
Each state
    %corresponds to a particular colour of fluorescence.
    figure(colours);
    red_t(fcount)=(red_i-red_f)*exp(-kcol*t(fcount))+red_f;
    subplot(3,N,fcount);
    set(gca,'Color',[1 1-red_t(fcount) 1-red_t(fcount)]);
    set(gca,'XTick',[],'YTick',[]);
    box on;
    green_t(fcount)=(green_i-green_f)*exp(-
kcol*t(fcount))+green_f;
    subplot(3,N,fcount+N);
    set(gca,'Color',[1-green_t(fcount) 1 1-green_t(fcount)]);
    set(gca,'XTick',[],'YTick',[]);
    box on;
    blue_t(fcount)=(blue_i-blue_f)*exp(-kcol*t(fcount))+blue_f;
    subplot(3,N,fcount+2*N);
    set(gca,'Color',[1-blue_t(fcount) 1-blue_t(fcount) 1]);
    set(gca,'XTick',[],'YTick',[]);
    box on;
end
```
